# Symbiosis modulates gene expression of symbionts, but not hosts, under thermal challenge

**DOI:** 10.1101/2023.11.21.568130

**Authors:** Hannah E Aichelman, Alexa K Huzar, Daniel M Wuitchik, Kathryn F Atherton, Rachel M Wright, Groves Dixon, E Schlatter, Nicole Haftel, Sarah W Davies

## Abstract

Increasing ocean temperatures are causing dysbiosis between coral hosts and their symbionts. Previous work suggests that coral host gene expression responds more strongly to environmental stress compared to their intracellular symbionts; however, the causes and consequences of this phenomenon remain untested. We hypothesized that symbionts are less responsive because hosts modulate symbiont environments to buffer stress. To test this hypothesis, we leveraged the facultative symbiosis between the scleractinian coral *Oculina arbuscula* and its symbiont *Breviolum psygmophilum* to characterize gene expression responses of both symbiotic partners *in* and *ex hospite* under thermal challenges. To characterize host and *in hospite* symbiont responses, symbiotic and aposymbiotic *O. arbuscula* were exposed to three treatments: 1) control (18°C), 2) heat (32°C), and 3) cold (6°C). This experiment was replicated with *B. psygmophilum* cultured from *O. arbuscula* to characterize *ex hospite* symbiont responses. Both thermal challenges elicited classic environmental stress responses (ESRs) in *O. arbuscula* regardless of symbiotic state, with hosts responding more strongly to cold challenge. Hosts also exhibited stronger responses than *in hospite* symbionts. *In* and *ex hospite B. psygmophilum* both downregulated genes associated with photosynthesis under thermal challenge; however, *ex hospite* symbionts exhibited greater gene expression plasticity and differential expression of genes associated with ESRs. Taken together, these findings suggest that *O. arbuscula* hosts may buffer environments of *B. psygmophilum* symbionts; however, we outline the future work needed to confirm this hypothesis.

## Introduction

Endosymbioses—associations where one organism lives within cells of its host [1]—have driven evolutionary innovations and allowed species to access resources and environments that would otherwise be unavailable [2, 3]. Endosymbioses span the tree of life and comprise exemplary innovations including deep-sea hydrothermal vent tubeworms (*Riftia pachyptila*) that rely on chemosynthetic bacterial endosymbionts [*e.g.*, 4] and salamanders (*Ambystoma maculatum*) benefiting from photosynthetic endosymbionts (*Oophila amblystomatis*) as embryos [*e.g.,* 5]. Endosymbionts often live within a host compartment, such as a vacuole or membrane, which facilitates the exchange of nutrients and metabolites, serving as the backbone for these symbioses [2, 6].

Corals are one of the most iconic examples of endosymbiosis, and their symbiosis with single celled algal symbionts [dinoflagellate algae in the family Symbiodiniaceae, hereafter ‘symbiont’, 7] enables diverse tropical reef ecosystems to thrive in oligotrophic waters [3]. Symbiodiniaceae live in coral gastrodermal cells in specialized vacuoles called symbiosomes [2, 8]. This endosymbiosis facilitates the transfer of materials between host and symbiont, where Symbiodiniaceae share photosynthetically-derived carbon sugars and in return receive inorganic compounds from coral metabolic waste in addition to protection [9, 10]. Once symbiosis is established, hosts can actively modulate symbiont physiology by manipulating the symbiosome environment. For example, symbiont photosynthesis is dependent on nitrogen availability, and host-mediated nitrogen limitation enables maintenance of primary production and control of symbiont growth [11, 12]. Additionally, coral hosts acidify the symbiosome *via* expression of V-type proton ATPases, which facilitates increased photosynthesis [13].

Tropical corals live near their upper thermal limits, making them particularly susceptible to temperature changes [14, 15]. Increases in anthropogenic carbon dioxide levels are elevating ocean temperatures and leading to marine heatwaves [*i.e.*, 16], which threaten corals globally [17]. Specifically, temperature increases lead to a breakdown of coral-algal symbioses in a process called ‘coral bleaching’, and extended periods of dysbiosis can lead to coral starvation and eventual mortality [18]. It is theorized that reactive oxygen species (ROS) generated by algal symbionts under temperature stress can damage cellular components, cause photoinhibition, and trigger coral bleaching [reviewed in 19]. However, even though both symbiotic partners exhibit a wide array of stress responses, symbionts are assumed to initiate dysbiosis due to ROS production [*e.g.,* 20–22]. Additionally, several lines of evidence demonstrate that hosts exhibit strong gene expression responses to stress [*e.g.,* 23–25, reviewed in 26], while the symbiont’s response is muted [*e.g.,* 27– 29]. This paucity of an algal response suggests that corals may regulate their symbiont’s environment to buffer algae from stress; however, alternative explanations include that the symbiont’s transcriptomic machinery is less responsive to stress regardless of symbiotic state.

Understanding the independent and interactive roles of coral hosts and Symbiodiniaceae algae in holobiont (*i.e.,* assemblage of coral host and associated algal and microbial symbionts) resilience is difficult in a tropical coral system [reviewed in 30] because it is impossible to disentangle the host’s aposymbiotic state from stress and nutrient limitation given tropical coral reliance on Symbiodiniaceae-derived carbon. To address these difficulties, facultative symbioses have emerged as tractable systems for answering fundamental questions about coral symbiosis [31]. Here, we used gene expression profiling in the facultatively symbiotic coral *Oculina arbuscula* and its symbiont *Breviolum psygmophilum* to address two questions: 1) what is the consequence of symbiosis for coral hosts under thermal challenge? and 2) how does symbiosis modulate symbiont responses to thermal challenges? We hypothesized that, compared to aposymbiotic corals, symbiotic corals under thermal challenge would exhibit gene expression patterns consistent with environmental stress responses (ESRs) of tropical corals because of symbiont-derived ROS produced under thermal stress. Based on previous work documenting minimal *in hospite* symbiont gene expression responses, we predicted greater responses of symbiotic hosts compared to symbionts *in hospite* and muted responses of symbionts *in hospite* compared to *ex hospite,* consistent with coral hosts modulating symbiont environments. To answer these questions, we conducted temperature challenge assays and characterized host and symbiont responses *via* gene expression profiling.

## Materials and Methods

### *I. Experiment 1:* Oculina arbuscula *and* Breviolum psygmophilum *holobiont responses to temperature challenges in symbiosis*

#### Coral Collection & Experimental Design

In June 2018, 16 *O. arbuscula* colonies (N=8 symbiotic, N=8 aposymbiotic) were collected from Radio Island, North Carolina (NC) (34.712590°N, -76.684308°W) under NC Division of Marine Fisheries permits #706481 and #1627488 (Figure 1A). Colonies were shipped overnight to Boston University, fragmented, attached to petri dishes using cyanoacrylate glue, and maintained at ambient conditions (18°C, 33-35 PSU) for approximately 5 months. Experimental temperatures were informed by 2017 (January 1 - December 31, 2017) *in situ* temperature data recorded by the NOAA buoy closest to the collection site (Station BFTN7; minimum temperature recorded=3°C, maximum temperature recorded=28.5°C; Figure 1B). One fragment from each colony (N=48 fragments) was placed in one of three treatments: 1) control: 18°C, 2) heat challenge: temperature increased 1°C day^-1^ from 18°C to 32°C, and 3) cold challenge: temperature decreased 1°C day^-1^ from 18°C to 6°C. Treatments were maintained for 15 days using Aqua Logic digital temperature controllers calibrated with a NIST-certified glass thermometer and were recorded using HOBO loggers (Figure 1C).

**Figure 1.**
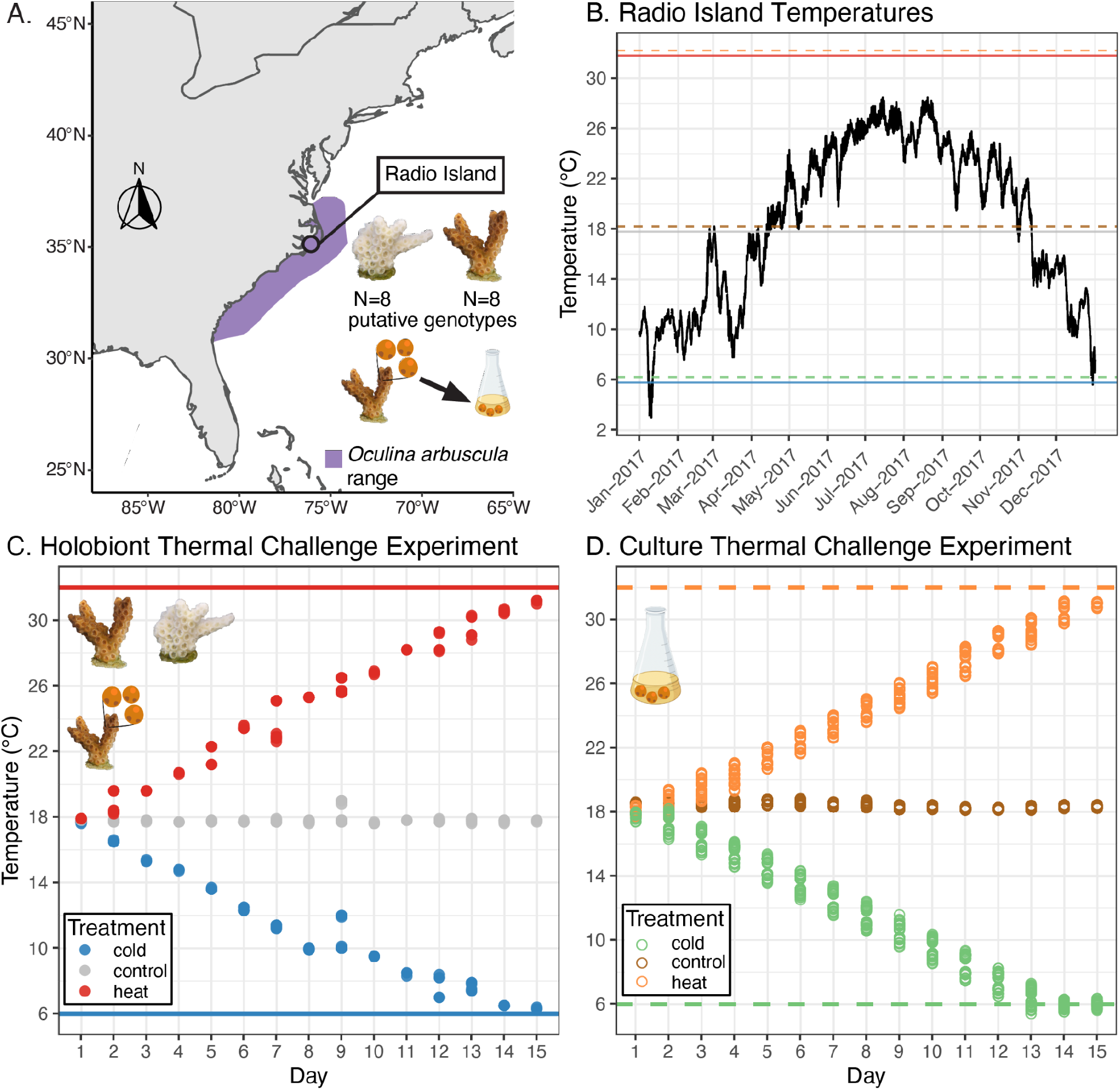
Experimental overview. **A.** Map of collection site (Radio Island, North Carolina) of *Oculina arbuscula* used in the holobiont thermal challenge experiment (N=8/symbiotic state). Purple shading indicates *O. arbuscula* range. **B.** Water temperatures (black line) in 2017 recorded at the NOAA buoy near Radio Island with thermal challenge treatments (heat=32°C, control=18°C, cold=6°C) overlaid. **C.** Temperatures recorded during holobiont thermal challenge experiment. **D.** Temperatures recorded during culture thermal challenge experiment. Culture icons were created with BioRender.com.

#### In hospite *symbiont physiology*

Pulse Amplitude Modulation (PAM) fluorometry measured dark-acclimated photochemical efficiency of photosystem II (Fv/Fm) of symbiotic corals using a Junior PAM approximately every three days throughout the experiment. Corals received 8 hours of dark acclimation before Fv/Fm was measured in triplicate. The effect of temperature challenge on symbiont Fv/Fm over time was analyzed using a linear mixed effects model (*lmer*), with interactions of fixed effects of treatment and day plus a random effect of genotype. Pairwise model outputs were compared using *emmeans* [32]. All analyses were performed in the R v4.2.0 statistical environment [33], and all code and raw data for the analyses detailed here can be found on Github: https://github.com/hannahaichelman/Oculina_Host_Sym_GE.

#### Oculina arbuscula *holobiont gene expression profiling*

On day 15, tissue from all fragments (N=48) was sampled using sterilized bone cutters, immediately preserved in 200 proof ethanol, and maintained at -80°C. Samples were homogenized with lysis buffer and glass beads and total RNA was extracted using an RNAqueous kit (ThermoFisher Scientific) following manufacturer’s instructions. DNA contamination was removed *via* DNAse1 and TagSeq libraries were prepared using 1.5 *μ*g of RNA [following 25, 34]. Successful libraries (N=47) were sequenced on Illumina HiSeq 2500 (single-end 50 bp) at Tufts University Core Facility. TagSeq analyses followed https://github.com/z0on/tag-based_RNAseq. Raw reads were quality filtered to remove Illumina adapters, poly-A sequences, PCR duplicates, reads less than 20 bp long, and reads with a quality score less than 33.

#### Putative coral clone identification

We tested for the presence of clones in our dataset following methods presented in Bove et al. [35]. Quality filtered reads were mapped to concatenated *O. arbuscula* and *B. psygmophilum* transcriptomes [presented in 36] using Bowtie2 [37]. Symbiont reads were removed from the dataset, genotyping and identification of host single nucleotide polymorphisms (SNPs) was performed using ANGSD [38], and putative clones were distinguished using a hierarchical clustering tree (*hclust*) based on pairwise identity by state (IBS) distances calculated in ANGSD (Figure S1). After removing clones, 33 samples (N=7 putative symbiotic genotypes, N=4 putative aposymbiotic genotypes) were used in downstream analyses. See supplementary methods for additional details.

#### *Oculina arbuscula* and *in hospite B. psygmophilum* gene expression analyses

Quality filtered reads were mapped to the same concatenated transcriptomes described above using Bowtie2 [37], but with different parameters (-k mode, k=5, --no-hd, --no-sq). Host and symbiont reads were separated to produce two separate files of counts per gene, and independent runs of DESeq2 [39] identified differentially expressed genes (DEGs) in response to heat and cold thermal challenge relative to control using Wald’s tests.

*Rlog*-transformed host and symbiont gene expression (GE) data were used as input for separate principal component analyses (PCAs) using *plotPCA* (package=DESeq2) to determine the effect of temperature on GE profiles using PERMANOVAs *via* the *adonis2* function [package=vegan; 40]. GE plasticity, defined as the distance in PC space between an individual’s GE profile and the average GE of all samples in the control, was calculated from host and symbiont PCAs following Bove et al. [35]. Differences in GE plasticity between treatments were tested using an ANOVA followed by Tukey’s HSD post-hoc tests. Model assumptions were assessed using *check_model* [package=performance; 41].

Gene ontology (GO) enrichment analyses were performed using Mann-Whitney *U* tests based on the ranking of signed log p-values [42] for both host and symbiont datasets. Results were visualized in dendrograms, which indicate the amount of gene sharing between significant GO categories and the direction of change relative to the control treatment. Results from the GO enrichment analyses were used for two functional analyses, detailed below.

First, GO delta-ranks, which quantify the tendency of genes assigned to a GO category toward up- or downregulation in treatment versus controls, were used to compare *O. arbuscula* host response under thermal challenges relative to a meta-analysis from Dixon et al. [26] that characterized the GE signatures of stress in *Acropora* corals. This meta-analysis identified two classes of coral stress responses: “type A”, which was positively correlated across projects and consistent with the coral environmental stress response (ESR), and “type B”, which was negatively correlated and indicated lower intensity stress. Host delta-ranks for Biological Processes (BP) GO terms were plotted against BP delta-ranks of all ‘type A’ studies identified by Dixon et al. [26]. While not a formal statistical test, this analysis indicates whether *O. arbuscula* responses were functionally similar to a ‘type A’ ESR. Second, *B. psygmophilum* symbiont GO results identified underrepresentation of photosynthesis-related GO terms under cold challenge. Genes associated with these GO categories (unadjusted p-value <0.10) were explored by constructing a heatmap using *pheatmap*.

Symbiont species identity of all symbiotic coral samples was confirmed using metabarcoding of the ITS2 locus (forward primer=*ITS-DINO* [43], reverse primer=*ITS2Rev2* [44]). Raw ITS2 data were submitted to SymPortal [45] to identify ITS2 defining intragenomic variant (DIV) profiles and relative abundances of DIVs across *O. arbuscula* fragments were compared using a bar plot constructed with *phyloseq* [46]. N=39/48 samples were successfully sequenced and confirmed to host *B. psygmophilum* (Figure S2). See supplementary methods for additional details on library preparation and sequencing.

#### *Comparing orthologous genes in* Oculina arbuscula *and* Breviolum psygmophilum *in symbiosis*

To compare *O. arbuscula* (symbiotic and aposymbiotic) and *B. psygmophilum in hospite* responses to temperature challenges, independent GE analyses were completed on orthologous genes. This analysis allowed us to test two predictions: 1) symbiotic hosts would respond more to temperature challenge than aposymbiotic hosts, and 2) symbiotic hosts would respond more than their symbionts *in hospite*. Orthologous genes were identified following Dixon and Kenkel [47] with additional specifics for Symbiodiniaceae described here: https://github.com/grovesdixon/symbiodinium_orthologs (details in supplemental methods). Briefly, Transdecoder v5.5.0 [48] predicted protein coding sequences, FastOrtho assigned predicted coding sequences to orthologous groups [49], and approximately-maximum-likelihood phylogenetic trees of these protein sequences were built using MAFFT [50] and FastTree [51], which resulted in a total of 1962 single-copy orthologs.

Single-copy orthologs (N=1962) were extracted from host and symbiont counts files and orthologs with mean count >2 in at least 80% of samples were retained, leaving 152 nonzero read count orthologs. To directly compare responses to thermal challenge of symbiotic hosts, aposymbiotic hosts, and symbionts *in hospite*, *O. arbuscula* and *B. psygmophilum* count data for these orthologs were analyzed in DESeq2, which modeled aggregate factors of temperature treatment and sample type (symbiotic host, aposymbiotic host, or symbiont *in hospite*). Responses of sample types to heat and cold challenge relative to control were quantified as the number of differentially expressed orthologs (DEOs, adjusted p-value <0.1). Differences in the proportion of DEOs across sample types were tested with a two-proportions z-test.

### *II. Experiment 2:* Breviolum psygmophilum *response in culture* - ex hospite

#### Symbiont cell culture maintenance

To isolate *B. psygmophilum* responses to temperature challenge *ex hospite*, *O. arbuscula* holobiont thermal challenges were replicated on cultured *B. psygmophilum*. *Breviolum psygmophilum* cells were isolated from *O. arbuscula* from Radio Island, NC by serially diluting airbrushed host tissue into sterile F/2 media (Bigelow NCMA, East Boothbay, ME, USA). The “ancestral culture” was maintained in F/2 media based on artificial seawater (Instant Ocean), with monthly transfers to fresh media, in a Percival incubator (model AL-30L2) at 26°C and irradiance of 30 *μ*mol photons m^-2^ sec^-1^ on a 14:10 hour light:dark cycle. After four months, this ancestral culture was split into three flasks, each with 100 mL of F/2 media and 0.5 mL of dense cells. These “daughter cultures” were acclimated to 18°C by decreasing temperatures at a rate of 1°C day^-1^ over a span of nine days. A preliminary experiment established semi-continuous culture methodology (Figure S3A; supplementary methods). Cultured symbiont species identity was confirmed prior to thermal challenge experiments using Sanger sequencing (details in supplementary methods).

#### Thermal challenge experiment

*Breviolum psygmophilum* cultures were exposed to thermal challenges that mirrored treatments used in holobiont experiments detailed in Part I. *Ex hospite* symbiont thermal challenges began after a 51-week acclimation at 18°C. Experimental cultures (N=4 flasks per treatment) were established from long-term acclimated daughter flasks, with initial cell densities of 200,000 cells mL^-1^ in 100 mL of F/2 media. Heat and cold challenge flasks were placed in separate Percival incubators (model AL-30L2), and control flasks were maintained in a temperature-controlled room (Harris Environmental Systems, Andover, MA). All treatments began at 18°C and followed a 14:10 hour light:dark cycle at ∼50 *μ*mol photons m^-2^ sec^-1^. Experimental cultures were grown semi-continuously, with the timing of transfers determined using the preliminary experiment (Figure S3).

#### Ex hospite Breviolum psygmophilum *gene expression profiling*

On day 15, all cultures were thoroughly mixed, concentrated *via* centrifugation (5000 RPM, 7 minutes), flash frozen in liquid nitrogen, and stored at -80°C. To obtain sufficient RNA for TagSeq, replicate cultures in cold challenge treatments were pooled, such that there were four pooled replicates extracted separately. Lower cell densities in cold challenge were due to reduced growth (Figure S3), which was not the case for heat challenge and control flasks. Flash frozen pellets were ground for three minutes in a mortar and pestle that was pre-chilled with liquid nitrogen. Additional liquid nitrogen was added to keep the cell pellet frozen and RNA was extracted using RNAqueous-micro kits (ThermoFisher Scientific) following manufacturer’s instructions, except elution volume was 15 *μ*L. DNA was removed *via* DNA-free DNA Removal Kit (ThermoFisher Scientific). RNA was normalized using concentrations from a Quant-iT PicoGreen dsDNA Assay (ThermoFisher Scientific) and total RNA was sent to the University of Texas at Austin Genome Sequencing and Analysis Facility (GSAF), where it was prepared for TagSeq following Meyer et al. [25] on a NovaSeq 6000 machine (single-end 100 bp).

#### *Gene expression analyses on* ex hospite Breviolum psygmophilum

Raw read processing of *ex hospite B. psygmophilum* TagSeq data followed methods detailed in Part I except samples were only mapped to the *B. psygmophilum* reference transcriptome. Principal component analysis (PCAs), GE plasticity, and GO-enrichment analyses were conducted as detailed in Part I. A heat-map of genes with GO annotations related to photosynthesis was generated on the culture dataset with an unadjusted p-value <0.01 to restrict the number of DEGs included. In addition, a heatmap of genes with GO annotations related to oxidative stress was constructed, as these terms were consistently enriched in *ex hospite* GO analyses.

### *III. Comparing* Breviolum psygmophilum *response* in *and* ex hospite

To test the prediction that symbionts would respond more to temperature challenge *ex hospite* compared to *in hospite*, *B. psygmophilum* GE datasets from Experiments I and II were analyzed together. A batch effect correction was conducted on combined raw count data using ComBat-seq [52], with a specified batch of experiment type (*in hospite* or *ex hospite*) and temperature treatment (heat challenge, cold challenge, or control) as the treatment of interest. Batch-corrected data were included in the same DESeq2 [39] model, which modeled a main effect of the aggregate factor of treatment (cold challenge, heat challenge, or control) and sample type (*in hospite* or *ex hospite*). Genes were only retained when present in at least 80% of samples (33/41 samples) at a mean count of 2 or higher, which left 1885 genes for downstream analyses.

Following PCAs detailed in Experiment I, combined symbiont count data were *rlog*-transformed and used as input for a PCA to test the effect of the aggregate factor of temperature treatment and sample type on GE. Significance was assessed with PERMANOVA, using the *adonis2* function [package=vegan; 40]. GE plasticity was also calculated following methods detailed above.

## Results

### *I. Independent responses of* Oculina arbuscula *and* Breviolum psygmophilum *to temperature challenges in symbiosis*

#### Oculina arbuscula *hosts exhibit stronger gene expression responses to cold challenge than heat challenge regardless of symbiotic state*

The aggregate factor of temperature treatment and symbiotic state had a significant effect on *O. arbuscula* host gene expression patterns (Figure 2A; *ADONIS p*=0.001) with cold challenge eliciting higher gene expression plasticity in both symbiotic (*Tukey HSD p*=0.013) and aposymbiotic (*Tukey HSD p*<0.001) hosts compared to heat challenge (Figure 2B; *p*<0.001). Symbiotic state did not influence gene expression plasticity within temperature treatments (cold challenge, *Tukey HSD p*=0.218; heat challenge, *Tukey HSD p*=0.992; Figure 2B).

**Figure 2.**
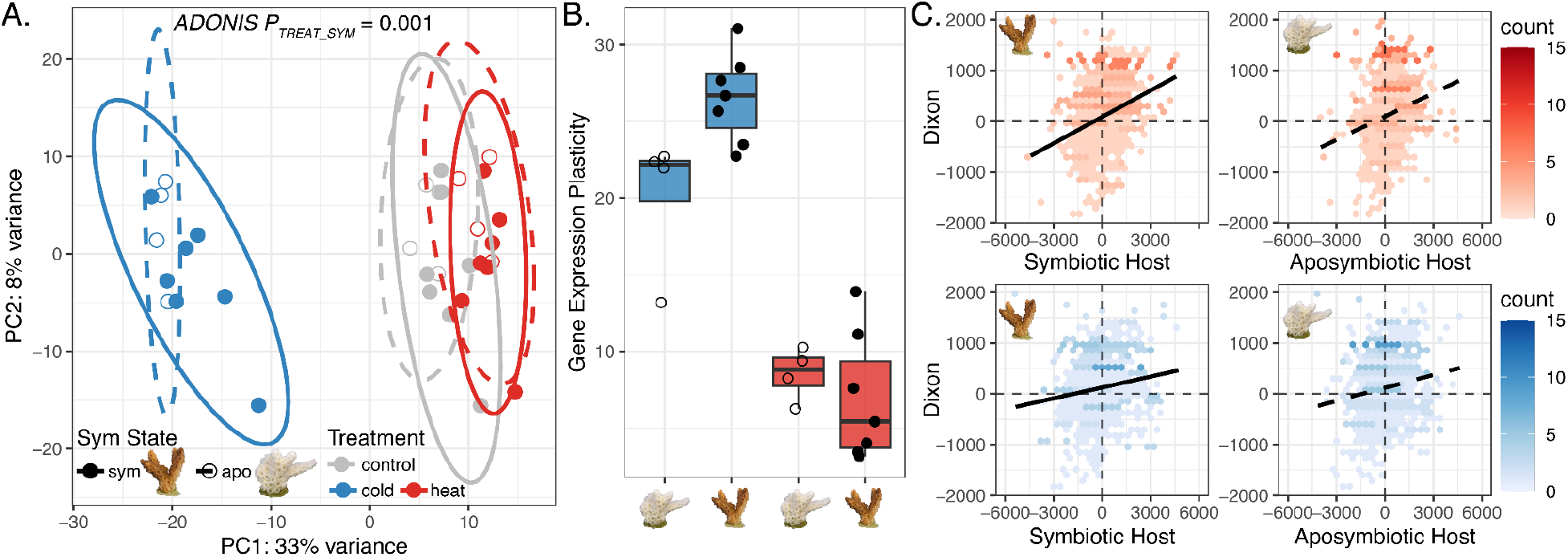
Symbiotic and aposymbiotic *Oculina arbuscula* host gene expression responses to temperature challenges. **A.** Principal component analysis (PCA) of gene expression of symbiotic (solid point and line) and aposymbiotic (open points, dashed line) coral hosts under control (grey), cold (blue), and heat (red) challenge. The x- and y-axes represent the % variance explained by the first and second PC, respectively. **B.** Gene expression plasticity of symbiotic and aposymbiotic hosts under cold and heat challenges. **C.** Relationship between biological processes (BP) gene ontology (GO) delta ranks from symbiotic (left) and aposymbiotic (right) hosts under heat (top) and cold (bottom) challenge with BP delta-ranks of all “type A” studies from [26]. Positive slopes represent “type A” environmental stress responses.

When comparing GO delta ranks of the ‘type A’ module from Dixon *et al.* [26] to symbiotic and aposymbiotic *O. arbuscula* host delta-ranks from the thermal challenges, positive relationships were observed for biological processes GO terms for all comparisons (Figure 2C).

#### *Negative effects of cold challenge on* Breviolum psygmophilum *photosynthetic function*

ITS2 metabarcoding confirmed all *O. arbuscula* genotypes hosted a majority of *B. psygmophilum* defining intragenomic variants (DIVs) (Figure S2). All but one individual hosted 100% *B. psygmophilum,* and all symbiotic *O. arbuscula* fragments hosted the same DIV of *B. psygmophilum* (Figure S2). Photosynthetic efficiency (Fv/Fm) of *in hospite B. psygmophilum* was reduced by temperature challenges through time (Figure 3A; *p*<0.001). By day 8 (cold challenge=11°C, heat challenge=25°C), Fv/Fm had significantly declined in the cold challenge relative to controls (*p*=0.02), but not in heat challenge (*p*=0.09). For the remainder of the experiment, Fv/Fm was significantly reduced in both cold and heat challenge relative to the control (Figure 3A; *p*<0.001 for all comparisons). Fv/Fm in cold challenge corals was more dramatically reduced than under heat challenge, with fixed effect parameter estimates on day 14 of -0.065 in heat challenge and -0.24 in cold challenge relative to control (Figure 3A).

**Figure 3.**
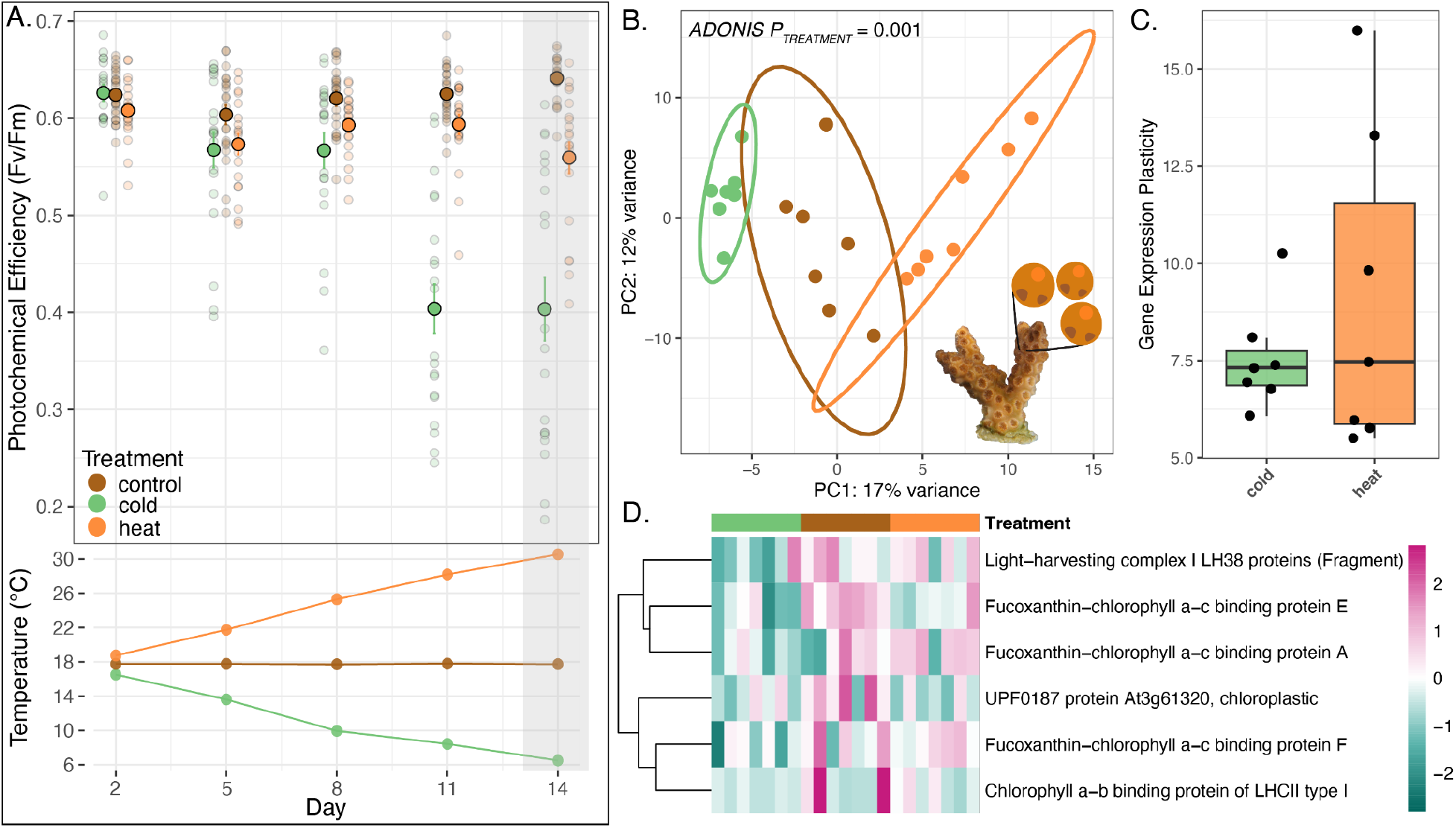
*In hospite* symbiont physiology and gene expression responses to temperature challenges. **A.** Photochemical efficiency (Fv/Fm, top panel) of *in hospite* symbionts through time as temperatures diverged (bottom panel). Top: Large points represent mean Fv/Fm ± standard error across treatments and transparent points represent a fragment’s Fv/Fm at each time point. The gray bar indicates the time point immediately prior to sampling for gene expression. **B.** Principal component analysis (PCA) of gene expression of *in hospite* symbionts from A on day 15. **C.** Gene expression plasticity of *in hospite* symbionts under cold (green) and heat (orange) challenge was not significantly different (Tukey HSD *p=*0.62). **D.** Heatmap showing differentially expressed genes (DEGs; unadjusted p-value<0.1) belonging to photosynthesis gene ontology (GO) terms, where each row is a gene and each column is a sample. The color scale represents log2 fold change relative to the gene’s mean, where pink represents up-regulation and teal represents down-regulation.

Temperature challenges significantly affected gene expression profiles of *in hospite B. psygmophilum* (Figure 3B; *ADONIS p=0.001*). However, no differences in gene expression plasticity between symbionts in cold and heat challenge were observed (Figure 3C; *Tukey HSD p*=0.62). Corroborating negative effects of cold challenge on *B. psygmophilum* Fv/Fm (Figure 3A), six GO terms related to photosynthetic processes were underrepresented in cold challenge relative to control conditions (photosystem [GO:0009521], photosynthesis, light harvesting [GO:0009765], chlorophyll binding [GO:0016168], protein-chromophore linkage [GO:0018298], thylakoid membrane [GO:0042651], and tetrapyrrole binding [GO:0046906]). Six annotated genes within these GO terms were downregulated under cold challenge (unadjusted p-value<0.10) relative to control conditions and these genes included light-harvesting complex (LHC) and chloroplast genes (Figure 3D).

### *II. Comparing response of* Oculina arbuscula *and* Breviolum psygmophilum *in symbiosis using orthologous genes*

More differentially expressed orthologs (DEOs) were observed in symbiotic hosts compared to aposymbiotic hosts under cold challenge (*X*^2^ = 4.76; *p*=0.015), but not heat challenge (Figure 4; *X*^2^ = 0.482; *p*=0.24). Similarly, aposymbiotic hosts exhibited more DEOs compared to *in hospite B. psygmophilum* under cold challenge (*X*^2^ = 37.7; *p*<0.0001), but not heat challenge (Figure 4; *X*^2^ = 0.781; *p*=0.188). The number of DEOs was greater in symbiotic hosts compared to *in hospite* symbionts under both cold challenge (*X*^2^ = 64.3; *p*<0.0001) and heat challenge (Figure 4; *X*^2^ = 3.23; *p*=0.036).

**Figure 4.**
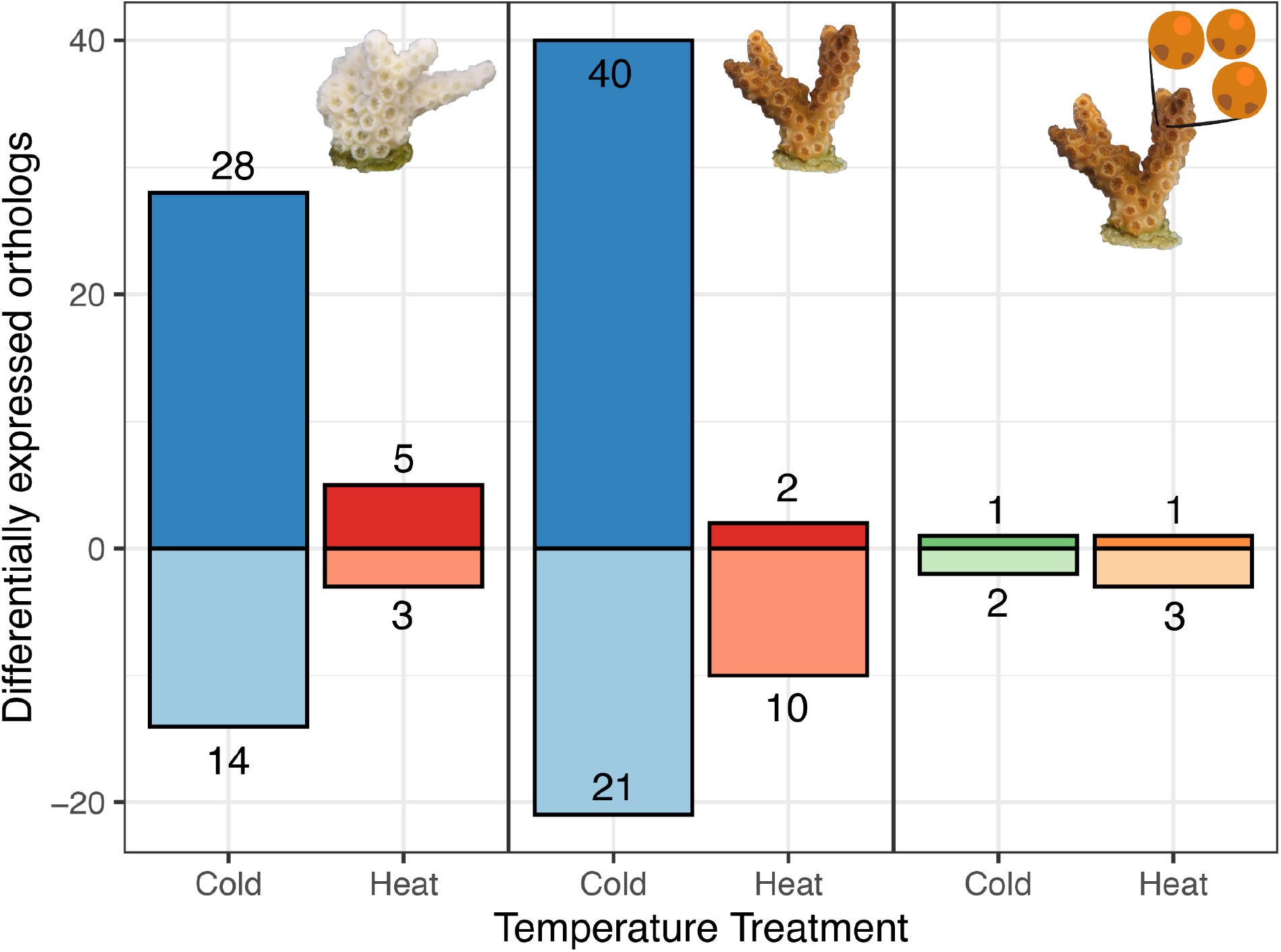
Coral hosts exhibit more differentially expressed orthologs (DEOs) than *in hospite* symbionts under thermal challenges. Bar plots representing the number of DEOs (positive values=up-regulated, negative values=down-regulated) in response to temperature challenges in aposymbiotic hosts (left), symbiotic hosts (center), and *in hospite* symbionts (right). Symbiotic *O. arbuscula* had significantly more DEOs than *in hospite B. psygmophilum* under cold challenge (*p<*0.0001) and heat challenge (*p=*0.036). Aposymbiotic *O. arbuscula* had significantly more DEOs than *in hospite B. psygmophilum* under cold challenge (*p<*0.0001), but not heat challenge (*p=*0.188). Symbiotic *O. arbuscula* had significantly more DEOs than aposymbiotic *O. arbuscula* under cold challenge (*p=*0.015), but not heat challenge (*p=*0.244).

### *III.* Breviolum psygmophilum *response to temperature challenge out of symbiosis* - ex hospite

Sanger sequencing confirmed that all parent cultures matched *B. psygmophilum* (GenBank Accession ID LK934671.1) with 100% percent identity and 53-87% query coverage. *Breviolum psygmophilum* cultures in all treatments were maintained in exponential growth phase throughout the experiment (Figure S3B,C). Thermal challenges significantly affected gene expression patterns of *ex hospite B. psygmophilum* (Figure 5A; *ADONIS p*=0.001). Gene expression plasticity was greater in *ex hospite B. psygmophilum* under cold challenge relative to heat challenge (Figure 5B; *Tukey HSD p*=0.008), whereas no significant effect was observed *in hospite* (Figure 3C).

Eight photosynthesis-related GO terms were underrepresented in *ex hospite B. psygmophilum* under cold challenge (photosystem [GO:0009521], photosynthesis, light harvesting [GO:0009765], chloroplast-nucleus signaling pathway [GO:0010019], photosynthesis [GO:0015979], chlorophyll binding [GO:0016168], protein-chromophore linkage [GO:0018298], thylakoid membrane [GO:0042651], and tetrapyrrole binding [GO:0046906]). A heat map of 59 DEGs (unadjusted p-value<0.01) under cold challenge belonging to these GO terms showcased a small group of up-regulated genes and a larger group of down-regulated genes under cold challenge (Figure 6A). Up-regulated photosynthesis-related genes included “Pentatricopeptide repeat-containing proteins”, which are involved in RNA editing events in chloroplasts [53]. Similar to *B. psygmophilum* in symbiosis (Figure 3C), genes involved in the LHC were down-regulated under cold challenge (Figure 6A).

**Figure 5.**
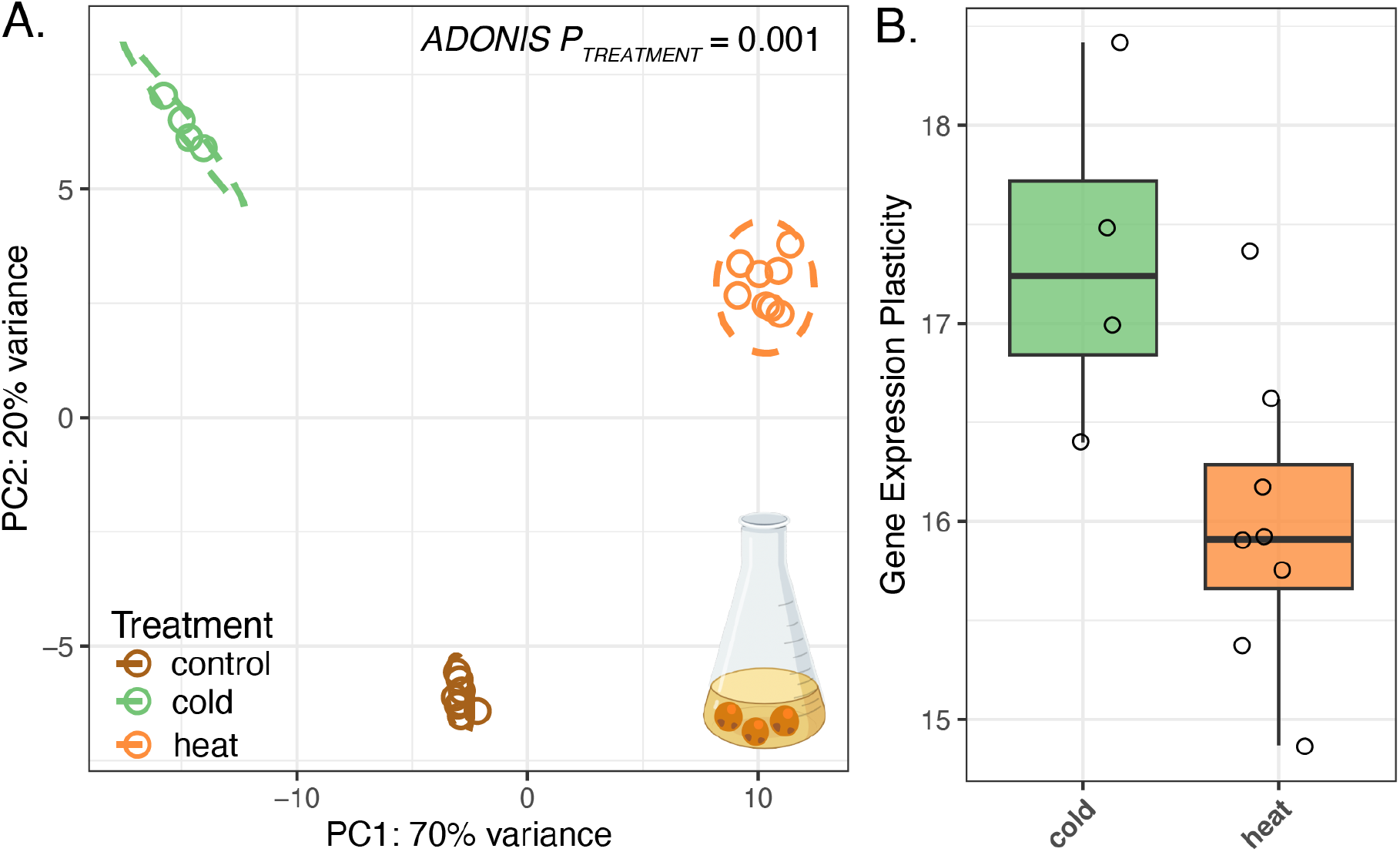
*Ex hospite* symbiont gene expression responses to temperature challenges. **A.** Principal component analysis (PCA) of gene expression of *ex hospite* symbionts under temperature challenges. **B.** Gene expression plasticity of symbionts *ex hospite* under cold (green) and heat (orange) challenge. Gene expression plasticity was significantly greater under cold challenge compared to heat challenge (*Tukey HSD p=*0.008). Culture icon created with BioRender.com.

**Figure 6.**
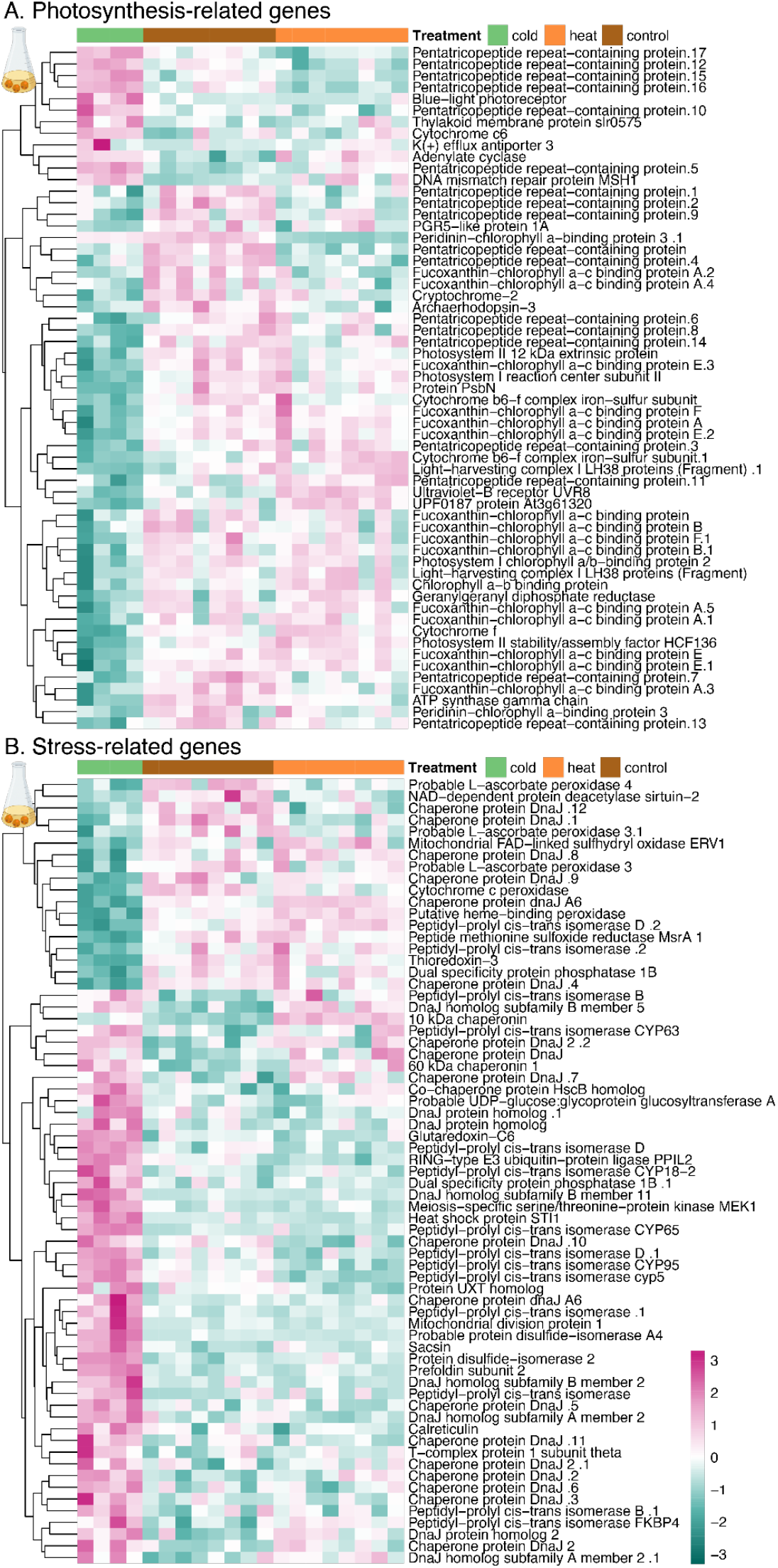
*Ex hospite* symbionts exhibit differential expression of photosynthesis and stress-related genes under cold challenge. Heatmap showing differentially expressed genes (DEGs; unadjusted p-value<0.01) belonging to photosynthesis (**A**) and stress (**B**) gene ontology (GO) terms, where each row is a gene and each column is a sample. The color scale represents log2 fold change relative to the gene’s mean, where pink represents up-regulation and teal represents down-regulation. Culture icons created with BioRender.com.

Five GO terms commonly associated with stress were differentially enriched in *ex hospite B. psygmophilum* under cold challenge relative to control conditions (protein folding [GO:0006457], cellular response to oxidative stress [GO:0034599], hydrogen peroxide metabolic process [GO:0042743], unfolded protein binding [GO:0051082], cellular response to chemical stress [GO:0062197]). A heat map of 69 DEGs under cold challenge (unadjusted p-value<0.01) assigned to these GO terms and revealed two groups of genes, one down-regulated and one up-regulated under cold challenge relative to control and heat challenge cultures (Figure 6B).

### *IV. Comparing responses of* in *and* ex hospite Breviolum psygmophilum

When analyzing *in hospite* and *ex hospite B. psygmophilum* in the same DESeq2 model, a significant effect of the aggregate factor of temperature treatment and symbiotic state was observed (Figure 7A; *ADONIS p*=0.001). Temperature and symbiotic state also had significant effects on gene expression plasticity (Figure 7B; *p<*0.0001), and *ex hospite B. psygmophilum* had significantly higher gene expression plasticity compared to *in hospite B. psygmophilum*, both under cold challenge (Tukey HSD *p*<0.0001) and heat challenge (Figure 7B; Tukey HSD *p*=0.0001).

**Figure 7.**
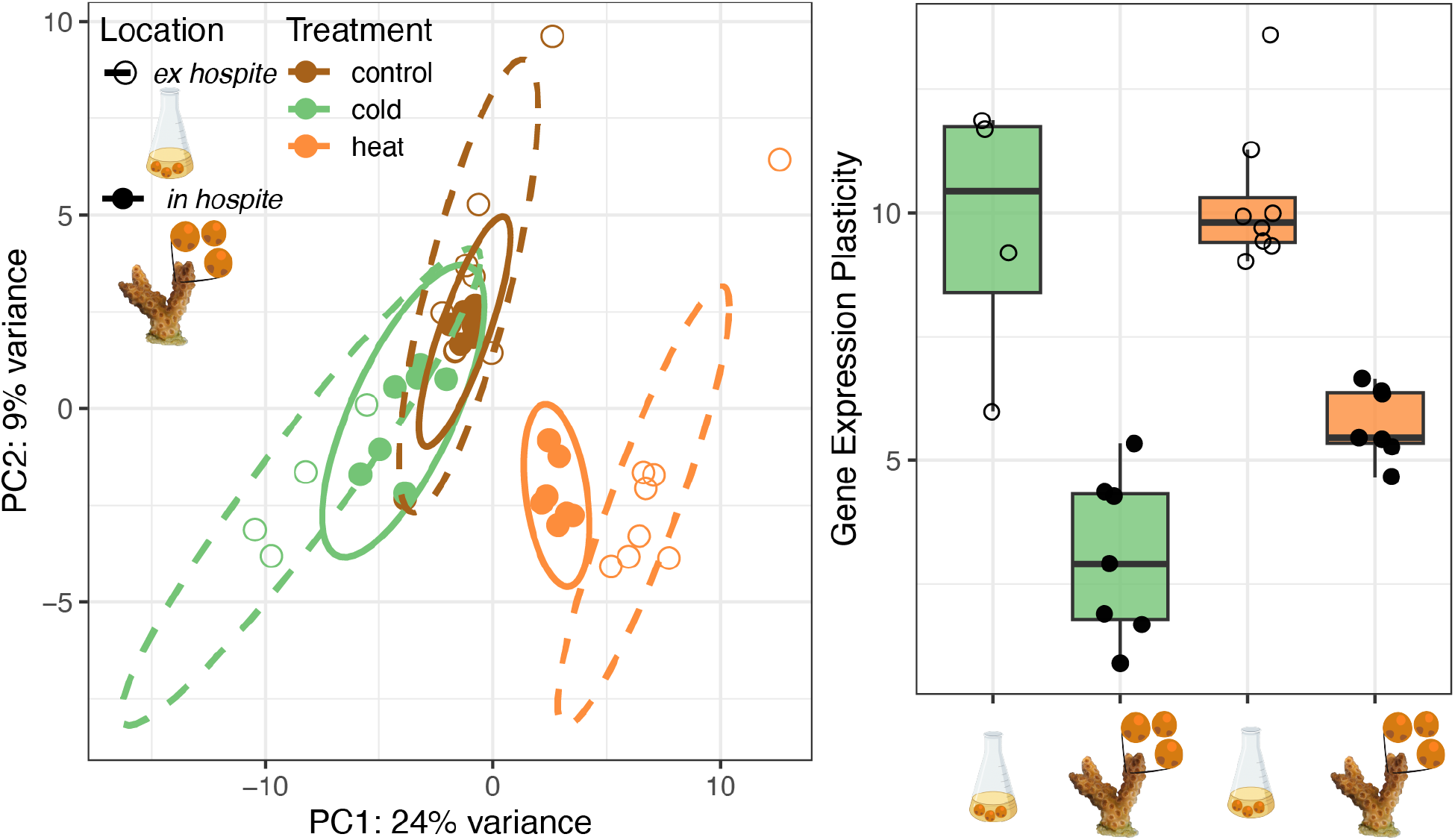
*Ex hospite* symbionts respond more strongly to thermal challenges than *in hospite* symbionts. **A.** Principal component analysis (PCA) of gene expression of *ex hospite* (open circles, dashed lines) and *in hospite* (solid points and lines) symbionts under control, cold, and heat temperatures. **B.** Gene expression plasticity of *ex hospite* and *in hospite* symbionts under cold (green) and heat (orange) challenge. Gene expression plasticity was significantly greater in *ex hospite* symbionts compared to *in hospite* symbionts under both cold challenge (Tukey HSD *p*<0.0001) and heat challenge (Tukey HSD *p*=0.0001). Culture icons created with BioRender.com.

## Discussion

### Both aposymbiotic and symbiotic coral hosts exhibit classic environmental stress responses to temperature challenges

We leveraged genome-wide gene expression profiling of *in* and *ex hospite* facultative coral hosts (*Oculina arbuscula*) and their algal symbionts (*Breviolum psygmophilum*) to disentangle the independent responses of hosts and symbionts to divergent thermal challenges across symbiotic states. In contrast to our prediction that symbiosis would alter the response of corals to thermal challenge, we found that both heat and cold challenges elicited general ESRs [‘type A’, 26] regardless of symbiotic state. Additionally, both symbiotic and aposymbiotic hosts exhibited greater gene expression plasticity in response to cold challenge compared to heat, aligning with previous work on the facultatively symbiotic coral *Astrangia poculata* when exposed to similar temperature challenges [54]. Wuitchik et al. [54] found that aposymbiotic *A. poculata* in cold challenge (6°C) exhibited five times more differentially expressed genes (DEGs) than in heat (31°C), corroborating the higher gene expression plasticity and number of differentially expressed orthologs (DEOs) observed in *O. arbuscula* under cold challenge. However, Wuitchik et al. [54] found cold challenge elicited a more severe ESR response [‘type A’, 26]) than heat challenge [‘type B’, 26], contrasting our findings that both thermal challenges elicited type A responses across symbiotic states. This pattern suggests that, even though *O. arbuscula* exhibited higher gene expression plasticity under cold challenge, corals in both temperature challenges were exhibiting stress responses consistent with a tropical coral’s ESR, highlighting the utility of *O. arbuscula* as a calcifying model for symbiosis [36].

The type A response presented in Dixon et al. [26] is characterized by functional enrichment of the coral ESR, including downregulation of cell division and upregulation of cell death, response to ROS, protein degradation, NF-*κ*B signaling, immune response, and protein folding. Specifically, type A datasets in tropical *Acropora* showcased upregulation of ROS and protein folding [26]. This informed our hypothesis that temperature challenge would result in an ESR-like response, akin to a type A response, in symbiotic *O. arbuscula* but not in aposymbiotic individuals. Instead, we observed that both symbiotic and aposymbiotic *O. arbuscula* exhibited type A responses under heat and cold challenge. This pattern could be the result of background symbionts in aposymbiotic corals [as previously observed in aposymbiotic *Astrangia poculata*, 55] producing ROS and leading to a type A response. Alternatively, aposymbiotic corals may be light-stressed as they lack shading from symbionts [*e.g.,* 56]. Additionally, the temperature challenges applied here were relatively short, and it is possible that symbiotic and aposymbiotic *O. arbuscula* would have exhibited differential responses if the challenges had been more extreme in temperature or duration [57]. In general, facultatively symbiotic corals are understudied, and future work should explore the responses of symbiotic and aposymbiotic corals under different stressors (*i.e.,* light, nutrients) and for longer durations [as in 58].

### *Evidence of host buffering in* O. arbuscula *holobionts*

We present three forms of evidence suggesting that *O. arbuscula* hosts are buffering their algal symbionts under thermal extremes: 1. The coral host exhibited more differentially expressed orthologs (DEOs) than its symbiont under cold challenge, 2. Stress-related genes were differentially expressed in symbionts *ex hospite* but not *in hospite*, and 3. *Ex hospite* symbionts exhibited higher gene expression plasticity in response to temperature challenges than *in hospite* symbionts. Higher gene expression plasticity in coral hosts compared to symbionts in symbiosis aligns with previous evidence suggesting that cnidarian hosts and their algal symbionts exhibit strong differences in the magnitude of gene expression responses under environmental challenges. For example, Davies et al. [59] reported that the tropical coral *Siderastrea siderea* exposed to 95-day temperature and ocean acidification challenges resulted in hosts consistently exhibiting greater differential expression of highly conserved genes compared to their symbiont *Cladocopium goreaui.* Barshis et al. [27] also found no changes in gene expression in either heat-susceptible *Cladocopium* (type C3K) or heat-tolerant *Durusdinium* (type D2) in symbiosis with *Acropora hyacinthus* following three days of high temperature exposure, which contrasted strong gene expression responses in the host [60]. Corroborating these patterns, Leggat et al. [29] observed that algae (*Cladocopium* C3) exhibited little change in expression of six stress and metabolic genes compared to their hosts (*Acropora aspera*).

Symbiosis itself has also been observed to alter gene expression patterns and physiology in Symbiodiniaceae algae. Here, we observed differential regulation of stress-related GO terms under cold challenge *ex hospite*, and genes within those terms included up-regulation of a heat shock protein (heat shock protein STI1) and a ubiquitin-related gene (RING-type E3 ubiquitin-protein ligase PPIL2). These genes are both classic ESR genes in tropical corals [26] and their differential regulation *ex hospite* highlights the potential benefits of a symbiotic lifestyle for Symbiodiniaceae. Symbiosis mitigating Symbiodiniaceae stress responses have been previously shown. For example, gene expression of *ex hospite Durusdinium trenchii* maintained at 28°C exhibited enrichment of the GO term “response to temperature stimulus” relative to *in hospite D. trenchii* in *Exaiptasia pallida,* which authors attributed to the protective microenvironment of the symbiosome [61]. Additionally, Maor-Landaw et al. [62] compared gene expression of *Breviolum minutum* in culture to *B. minutum* freshly isolated from *Exaiptasia diaphana* and observed down-regulation of genes indicative of the protected and stress-reduced environment of the symbiosome.

Specifically, pentatricopeptide repeats (PPR), which have been previously associated with Symbiodiniaceae RNA processing in response to environmental stress and were included in the repertoire of “stress responsive genes’’ in *Fugacium kawautii* [63], were down-regulated in freshly isolated *B. minutum* [62]. These findings support our third piece of evidence for host buffering. Together, these data suggest that *in hospite* symbionts respond less at the level of gene expression to cope with temperature challenges compared to *ex hospite* symbionts, providing further evidence that cnidarian hosts exert control over the symbiont’s micro-environment under environmental stress.

### *Cold challenge elicited negative effects on photosynthesis of* ex hospite *and* in hospite *Breviolum psygmophilum*

Although responses of *B. psygmophilum in hospite* were muted (*i.e.,* fewer DEGs and DEOs) under temperature challenges compared to responses *ex hospite,* negative effects on photosynthesis were observed at the level of phenotype (*in hospite* Fv/Fm) and gene expression (both *in hospite* and *ex hospite*), particularly under cold challenge. *Ex hospite B. psygmophilum* exhibited differential expression of photosynthesis and stress genes, including down-regulation of light harvesting complex (LHC) genes under cold challenge. This aligns with previous work investigating how symbiosis affects Symbiodiniaceae photosynthesis. For example, Bellantuono et al. [61] found that photosynthetic processes were modified in *D. trenchii* living *in hospite* compared to *ex hospite.* Specifically, GO terms related to photosynthesis (*i.e.*, photosynthesis, photosystem II repair, and light harvesting) were enriched *in hospite* compared to *ex hospite*, which may result from host carbon concentrating mechanisms increasing the availability of CO_2_ *in hospite*. In addition, we observed reduced Fv/Fm of *in hospite B. psygmophilum*, aligning with previous work demonstrating reduced Fv/Fm in cultured *B. psygmophilum* exposed to simulated seasonal temperature declines (cooled from 26°C to 10°C and for two weeks before returning to 26°C) [64]. In that study, *B. psygmophilum* Fv/Fm recovered to pre-challenge values once temperatures were returned to baseline, while other Symbiodiniaceae species that typically associate with tropical corals failed to regain Fv/Fm following cold challenge [64]. This difference was attributed to *B. psygmophilum’s* symbiosis with corals in temperate/subtropical areas where they experience larger annual temperature variation, aligning with a recent report of its wide thermal breadth (16.15°C) compared to six other Symbiodiniaceae isolates [65]. Therefore, Fv/Fm declines coupled with down-regulation of photosynthesis genes could represent *B. psygmophilum*’s seasonal response to low temperatures, and if the cultures were returned to control conditions, they may have recovered.

While we only observed strong phenotypic and gene expression responses of *B. psygmophilum* under cold challenge and not heat challenge, these links between photosynthetic disruption and transcriptional regulation of photosynthetic machinery align closely with previous work on Symbiodiniaceae under heat stress. Previous work has highlighted that heat stress can inhibit the synthesis and resulting mRNA pool of an antenna protein of the light harvesting complex (acpPC) in Symbiodiniaceae [66]. Additionally, temperature anomalies can alter thylakoid membrane fluidity, decoupling light harvesting and photochemistry, which suppresses ATP synthesis and results in increased reactive oxygen species (ROS) in Symbiodiniaceae [67]. LHCs are well-known for transferring absorbed light energy to the photosynthetic reaction center, but they also play an important role in photoprotection and have been linked to thermal sensitivity in Symbiodiniaceae [66]. Decreasing the number of peripheral LHCs may serve as a photoprotective mechanism under heat stress, as it ultimately decreases the light reaching photosynthetic reaction centers and reduces the risk of damage to D1 reaction center proteins [68]. It is possible that LHC down-regulation may represent a photoprotective mechanism in *B. psygmophilum* under any thermal stress. While Symbiodiniaceae gene expression in response to cold challenge remains largely unexplored, evidence suggests that cold challenge induces similar photophysiology responses as heat challenge in Symbiodiniaceae [*e.g.,* 64, 69–72]. It is therefore possible that the heat challenge explored here was not high enough or long enough to elicit a similarly negative response as cold challenge. Indeed, Fv/Fm of *in hospite B. psygmophilum* under heat challenge was declining, but remained significantly higher than in cold challenge. This pattern aligns with Roth et al. [72], who found that cold challenge was more immediately harmful for *Acropora yongei* symbiosis, but heat stress was more harmful in the long term. Although *ex hospite B. psygmophilum* Fv/Fm was not quantified here, differential regulation of photosynthesis and stress-related genes suggests an even stronger negative impact of cold stress on photosynthesis *ex hospite.* Altogether, future work would benefit from longer and more extreme temperature challenges to ensure that the entire thermal niche is investigated. Lastly, future studies quantifying additional phenotypes in both *in* and *ex hospite B. psygmophilum* to determine if such convergent responses to temperature challenge exist are warranted.

### Alternative hypotheses for “host buffering” and proposed future experiments

Transcriptional regulation is only one molecular process involved in responding to thermal stress. Mounting evidence suggests that a lack of transcriptional response under environmental challenges could be the result of post-transcriptional and/or post-translational mechanisms in Symbiodiniaceae. This includes evidence of microRNA (miRNA)-based gene regulatory mechanisms in Symbiodiniaceae [28, 73], which aligns with the genomic evidence that dinoflagellates may be more capable of translational rather than transcriptional regulation [74]. Previous work in *Symbiodinium microadriaticum* highlighted that a lack of common transcription factors and few DEGs could be attributed to small RNA (smRNA) post-transcriptional gene regulatory mechanisms [28]. It has also been proposed that the lack of transcriptional regulation in Symbiodiniaceae could be due to gene duplication as a mechanism to increase transcript and protein levels of genes [75]. It is therefore possible that the lack of *in hospite* symbiont response to thermal challenges found here is not evidence of ‘host buffering’, but instead that other post-transcriptional or post-translational regulation occurs more commonly in Symbiodiniaceae *in hospite*. Another unique aspect of Symbiodiniaceae genomes is *trans-*splicing of spliced leader sequences, which converts polycistronic mRNAs (code for multiple proteins) into monocistronic mRNAs (code for one protein) and potentially regulates gene expression [76, 77]. TagSeq cannot account for splice variation [25], preventing us from considering these differences. Additionally, lower depth of coverage of *in hospite B. psygmophilum* sequencing data captured here could have limited our ability to detect transcriptional responses. Finally, comparing algae in culture to algae in symbiosis inherently includes a confounding variable of nutritional status, as algae in culture exist in nutrient replete conditions [78]. Below we highlight future studies and data needed to further test the host buffering hypothesis.

Our work supports a scenario in which coral hosts modulate the environment of *in hospite* Symbiodiniaceae algae to buffer their responses to temperature challenges; however, additional experiments are needed to validate this hypothesis. First, replicating the temperature challenges performed here but leveraging proteomic and gene expression profiling in parallel [*e.g.*, 79] will be critical in establishing whether lack of gene expression of *in hospite* symbionts translates to a lack of proteomic response under stress. Including nutrient controls, namely *ex hospite* Symbiodiniaceae in nutrient-depleted media that replicate the nutritional environment of the symbiosome, are necessary to address the confounding variable of nutritional status. Additionally, this proposed experiment should implement RNA extraction methods that prioritize obtaining and sequencing equal amounts of genetic material from host and symbiont.

While understanding the response of subtropical corals to thermal extremes is valuable in its own right, the facultative symbiosis, calcifying nature, and available genomic resources of *O. arbuscula* make it a unique model for linking results to tropical coral responses as climate change progresses [36]. If coral hosts are able to regulate the environments of their symbionts, and this regulation in turn can serve to limit stress in the holobiont and ultimately reduce coral bleaching, then this host buffering phenotype may allow for the identification of coral-algal pairings that will be more resilient under future global change conditions.

## Acknowledgements

We would like to thank Steve Broadhurst for assistance collecting *Oculina arbuscula* colonies and Dr. Joel Fodrie for assistance with collection permits. The symbiont shapes used in figures were created by Giulia Puntin and modified for use (https://github.com/sPuntinG/Coral_stuff). Thank you to Carlos Tramonte for the *Oculina arbuscula* photos used in figures, as well as Alexandra Mercurio, Olivia Aswad, Julia Russo, Justin Scace, Dr. Nicola Kriefall, Brooke Benson, Julia Mendez, and the Boston University Marine Program (BUMP) for support and assistance with the coral holobiont experiment. Thank you to the BU Shared Computing Cluster for computational resources used to complete analyses. Finally, thank you to Dr. Peter Buston, Dr. Karen Warkentin, Dr. Sean Mullen, and Dr. Carly Kenkel for their thoughtful feedback on this manuscript.

## Data Accessibility and Benefit-Sharing Section

## Data Accessibility Statement

Raw sequencing data will be deposited in the SRA upon publication (BioProject ID=XX). All other data, code, and materials used in the analyses can be found on the Github repository associated with this project (https://github.com/hannahaichelman/Oculina_Host_Sym_GE) and additionally will be hosted on Zenodo upon publication (doi: X).

## Author Contributions

HEA and SWD designed research; HEA, AKH, DMW, KFA, ES, NH, GD, and RMW performed research; HEA and GD analyzed data; HEA and SWD wrote the paper. All co-authors provided final feedback on the manuscript.

