## Supplemental file for "Symbiosis modulates gene expression of symbionts, but not hosts, under thermal challenge"

**Supplemental Materials for "Symbiosis modulates gene expression of symbionts, but not hosts, under thermal challenge".**

**Authors:** Hannah E Aichelman[1*], Alexa K Huzar[1], Daniel M Wuitchik[1], Kathryn F Atherton[1], Rachel M Wright[1,2], Groves Dixon[3], E Schlatter[1], Nicole Haftel[1], and Sarah W Davies[1*]

### Supplementary Methods

#### *I. Experiment 1.* Oculina arbuscula *and* Breviolum psygmophilum *holobiont responses to temperature challenges in symbiosis*

*Coral Collection & Experimental Design*

Each temperature treatment consisted of three 15-gallon aquaria connected to one sump. All aquaria had a powerhead for water circulation, and each sump was equipped with a filter sock and protein skimmer for filtration. Water quality was tested daily in each tank by measuring temperature using a NIST-calibrated thermometer (Figure 1C) and salinity with a YSI meter. Target salinity of 33-34 PSU was maintained by mixing DI water with Instant Ocean Sea Salt, and mean (± SE) salinity was 33.94 ± 0.025 in control, 33.87 ± 0.015 in cold challenge, 34.15 ± 0.024 in heat challenge. A 50% water change was performed on day nine. Light exposure was monitored to ensure corals received equal light (50 µmol photons $m^2$ $sec^{-1}$) and remained on a 12:12 hour, light:dark schedule throughout the experiment. Coral fragments were rotated daily to ensure even light exposure. Each aquaria was fed ¼ tsp of reconstituted powdered brine shrimp daily, and feeding occurred for one hour before recirculating flow was resumed.

*Putative coral clone identification*

Quality filtered reads were mapped to concatenated *O. arbuscula* and *B. psygmophilum* transcriptomes [1] using Bowtie2 [2] with local mode (--local), seed substring alignment length of

16 (-L 16), suppressing records for unaligned reads (--no-unal), and a minimum alignment score function of f(x) = 16 + x, where x is read length (--score-min L,16,1). Symbiont reads were then removed from the dataset, and genotyping and identification of host SNPs was performed using ANGSD [3]. Loci were filtered to include those that were present in at least 80% of individuals, with a minimum mapping score of 20, a minimum quality score of 25, a strand bias p-value >1 x $10^{-5}$, a heterozygosity bias >1 x $10^{-5}$, a minimum minor allele frequency >0.05, a p-value >1 x $10^{-5}$, all triallelic sites were excluded as well as those with multiple best hits. Putative clones were distinguished using a hierarchical clustering tree (*hclust*) based on pairwise identity by state (IBS) distances calculated in ANGSD (Figure S1A). One genotype (A) was removed from downstream analyses because its replicate fragments failed to cluster together, suggesting sequencing failure, and the hierarchical clustering tree was re-made without that genotype (Figure S1B). This tree identified three sets of putative clones: 1) aposymbiotic putative clonal group of genotypes N, O, and P, 2) aposymbiotic putative clonal group of genotypes H and K, and 3) symbiotic putative clonal group of M and L (Figure S1B). The genotype within each putative clonal group with the highest depth of coverage was maintained in downstream analyses (N, M, and H) and all others were removed leaving a total of 33 samples (N=7 putative symbiotic genotypes, N=4 putative aposymbiotic genotypes).

*Confirming identity of* Breviolum psygmophilum in hospite

DNA was extracted from N=39/48 coral fragments that had sufficient sample remaining using a modified phenol-chloroform extraction, described in detail by Davies et al. [4]. Because aposymbiotic corals can still host a small amount of symbionts, both symbiotic and aposymbiotic fragments were included in these extractions. The ITS2 region was targeted using forward primer

*ITS-DINO* (5' - TCGTCGGCAGCGTC AGATGTGTATAAGAGACAG *NNNN* **GTGAATTGCAGAACTCCGTG** - 3') [5] and reverse primer *ITS2Rev2* (5' - GTCTCGTGGGCTCGG AGATGTGTATAAGAGACAG *NNNN* **CCTCCGCTTACTTATAGCTT** 3') [6]. Underlined bases denote adapter linker, bold bases are primer sequences, and the middle bases are spacer sequences. The reactions totaled 20 µl and included 20 ng of template DNA, 10 µM forward primer, 10 µM reverse primer, 0.2 mM dNTP, 1X ExTaq buffer (Takara), 0.025 U ExTaq enzyme (Takara), and the remaining Milli-Q $H_2O$ (Millipore). The PCR profile was 95°C for 40 seconds, 59°C for 120 seconds, and 72°C for 60 seconds for 35 cycles with a final elongation step of 72°C for 7 minutes. PCR products were purified using Ampure XP Reagent for PCR Purification (Beckman Coulter) and eluted in 28 µL. Each PCR product was barcoded with a unique Illumina barcoded adapter using five PCR cycles and visualized on a 1% agarose gel to assess relative band intensity. Samples were normalized and pooled and 25 µl of the pooled library was run on a 1% SYBR Green (Invitrogen) stained gel. The target band was excised and incubated with 30 µl of Milli-Q water overnight at 4˚C. This library was quantified using a Quant-iT PicoGreen dsDNA assay kit (Thermo Fisher) and submitted for paired-end 250 bp sequencing on an Illumina MiSeq at Tufts University Core Facility (TUCF).

*Comparing orthologous genes in* Oculina arbuscula *and* Breviolum psygmophilum *in symbiosis*

Orthologous genes were identified following Dixon and Kenkel [7] with additional specifics for Symbiodiniaceae described here: https://github.com/grovesdixon/symbiodinium_orthologs. Cd-hit [8] clustered sequences in *O. arbuscula* and *B. psygmophilum* reference transcriptomes with a sequence identity threshold of 0.98, alignment coverage of the longer and shorter sequence of at least 0.3, and only the longest sequence was retained. Transdecoder v5.5.0 [9] predicted protein coding sequences in the clustered references based on open reading frames (ORFs) and homology to known proteins. Only the longest ORFs (minimum amino acid length=50 bp) were retained and annotated using a blastp alignment against the Swissprot database, and protein domains were identified with scanHmm in HMMER v3.2.1 [10]. FastOrtho assigned these predicted coding sequences to orthologous groups with an e-value cut-off of $1 \times 10^{-10}$ [11]. Paralogs (N=9727 groups) were removed, leaving 1951 orthologous groups. Protein sequences for these orthologs were aligned using MAFFT [12] and gene trees were built with FastTree [13], which infers approximately-maximum-likelihood phylogenetic trees from protein sequences. These constructed trees were pruned using the biopython module *Phylo* [14], which facilitated the inclusion of additional orthologous groups as single copy orthologs, for a total of 1962 single-copy orthologs.

***II. Experiment 2.*** **Breviolum psygmophilum *response in culture* - ex hospite**

*Semi-continuous culture methodology*

After the daughter cultures reached 18°C, they were acclimated to 18°C for 11 weeks, after which new “test cultures” were created from the daughter cultures (one from each), which were used in a preliminary experiment to determine the timing of when cultures reached the stationary growth phase (Figure S3A). This information was leveraged to maintain cultures in exponential growth phase throughout the temperature challenge experiments. Each test culture initially had a cell density of 10,000 cells $mL^{-1}$ in a total volume of 100 mL F/2 media. Test cultures were maintained at 18°C under a 14:10 hour light:dark cycle when determining timing of exponential and stationary growth phases. Triplicate hemocytometer cell counts were conducted every other day on each flask and were used to calculate cell densities to establish timing of when the stationary growth phase was reached, which was approximately ten days after initial transfer (Fig S3A).

*Checking identity of* Breviolum psygmophilum ex hospite

*Ex hospite* symbiont species identity was confirmed prior to thermal challenge experiments. Daughter cultures were sub-sampled one week before the start of the experiment, on October 23, 2020. DNA was extracted using the DNeasy Plant Mini Kit (Qiagen) following manufacturer's instructions. The ITS2 region was targeted using the same forward and reverse primers and PCR profiles described above: *ITS-DINO* [5] and *ITS2Rev2* [6]. Amplified samples were sent to Eton Biosciences, where they were purified and sequenced using Sanger sequencing. Sequence quality was checked using *4Peaks*, and sequence identity was confirmed using NCBI Nucleotide BLAST with default parameters.

*Thermal challenge experiment*

Temperatures were changed in the heat and cold challenge treatments approximately 20 hours after the experimental cultures were created. Each temperature increase in successive days occurred during the dark phase of the light cycle at the same time each day (11:30). All experimental cultures were sub-sampled every other day for hemocytometer counts to track cell growth through time. On day 8, all cultures were homogenized, half of their volume (50 mL) was transferred to a sterile flask, and an equivalent volume of F/2 media was added. This doubled the number of experimental flasks, from N=12 to N=24 (N=8 replicate flasks per treatment). The 15-day culture experiment mirrored the holobiont experiment and final sample collection and processing was completed on November 13, 2020.

### Supplemental Figures

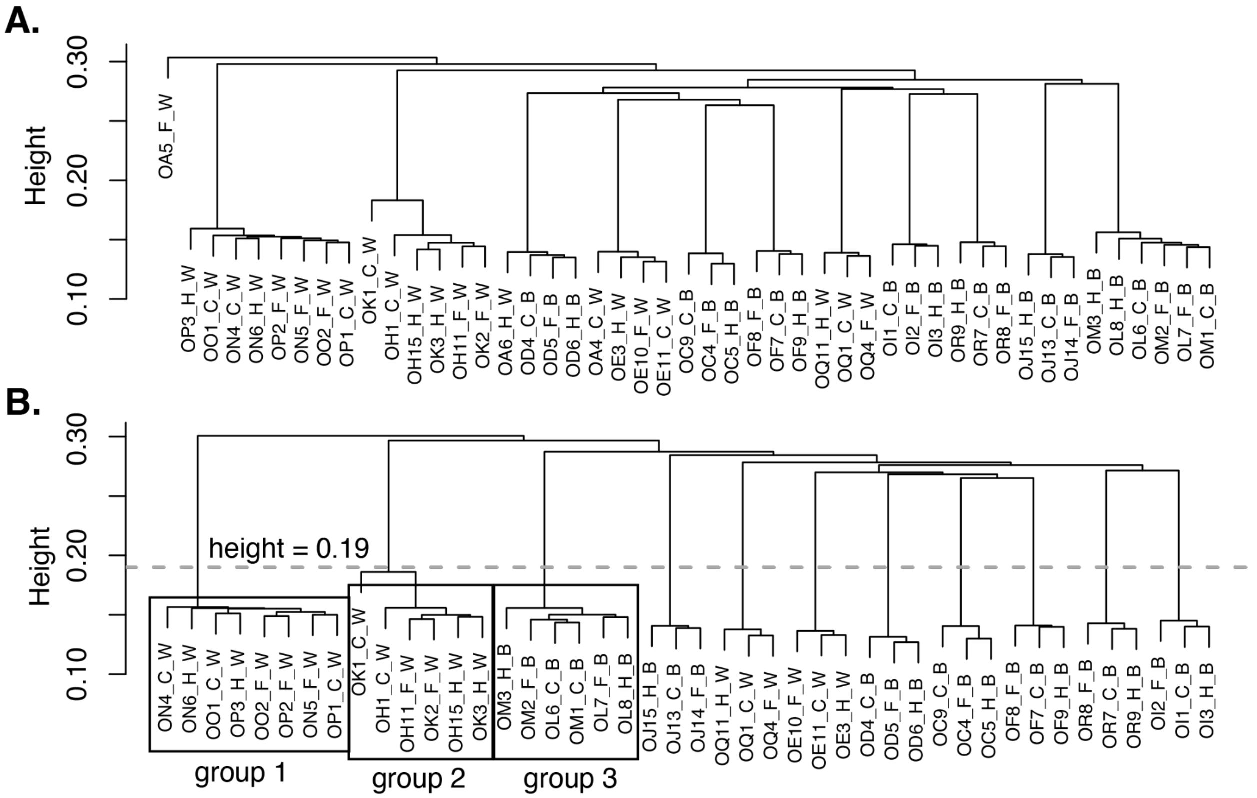


**Figure S1.** Identity by state (IBS) cluster dendrograms of *Oculina arbuscula* fragments, both with (**A**) and without (**B**) genotype A. (**B**) IBS cluster dendrogram indicates three putative clonal groups each containing more than one putative genotype, identified with black boxes. The dashed line at height=0.19 represents the cutoff for clone assignments, and the remaining groups outside of the three boxes indicate replicate fragments from the same genotype distributed across temperature challenge treatments. Sample IDs include information on genotype, fragment number, temperature challenge treatment, and symbiotic state. All samples start with 'O', indicating the coral species *O. arbuscula*, followed by a second letter denoting genet and a number, which denotes the fragment number. The letter following the first '_' indicates temperature treatment (C=control, F=cold, H=heat) and the final letter indicates symbiotic state (W=aposymbiotic (white), B=symbiotic (brown)).

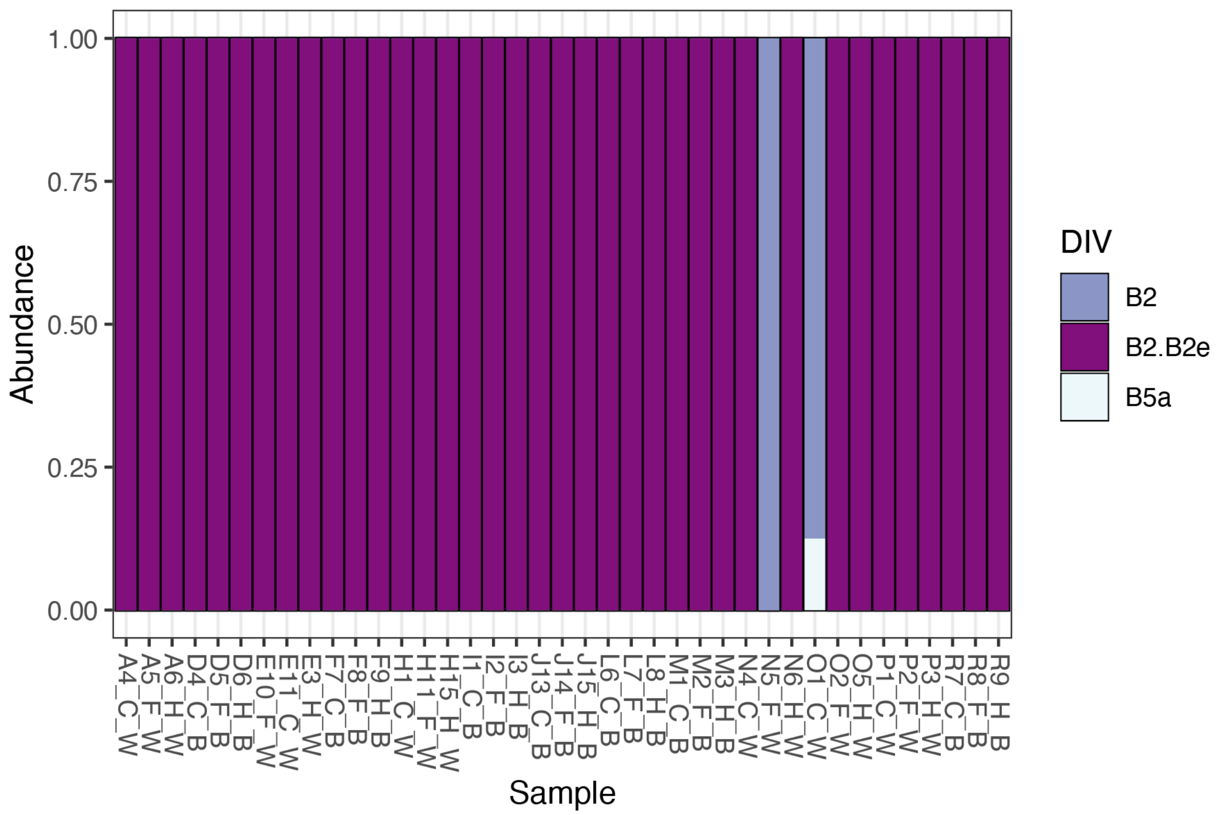


**Figure S2.** Bar plots of Symbiodiniaceae defining intragenomic variants (DIVs) hosted by *Oculina arbuscula*, colored by DIV. Each column of the bar plot represents one *O. arbuscula* fragment. Sample names include genotype, fragment number, temperature treatment (C=control, F=cold challenge, H=heat challenge), and symbiotic state (W=aposymbiotic (white), B=symbiotic (brown)). For example, A4_C_W can be interpreted as follows: genotype A, fragment 4, control treatment, aposymbiotic.

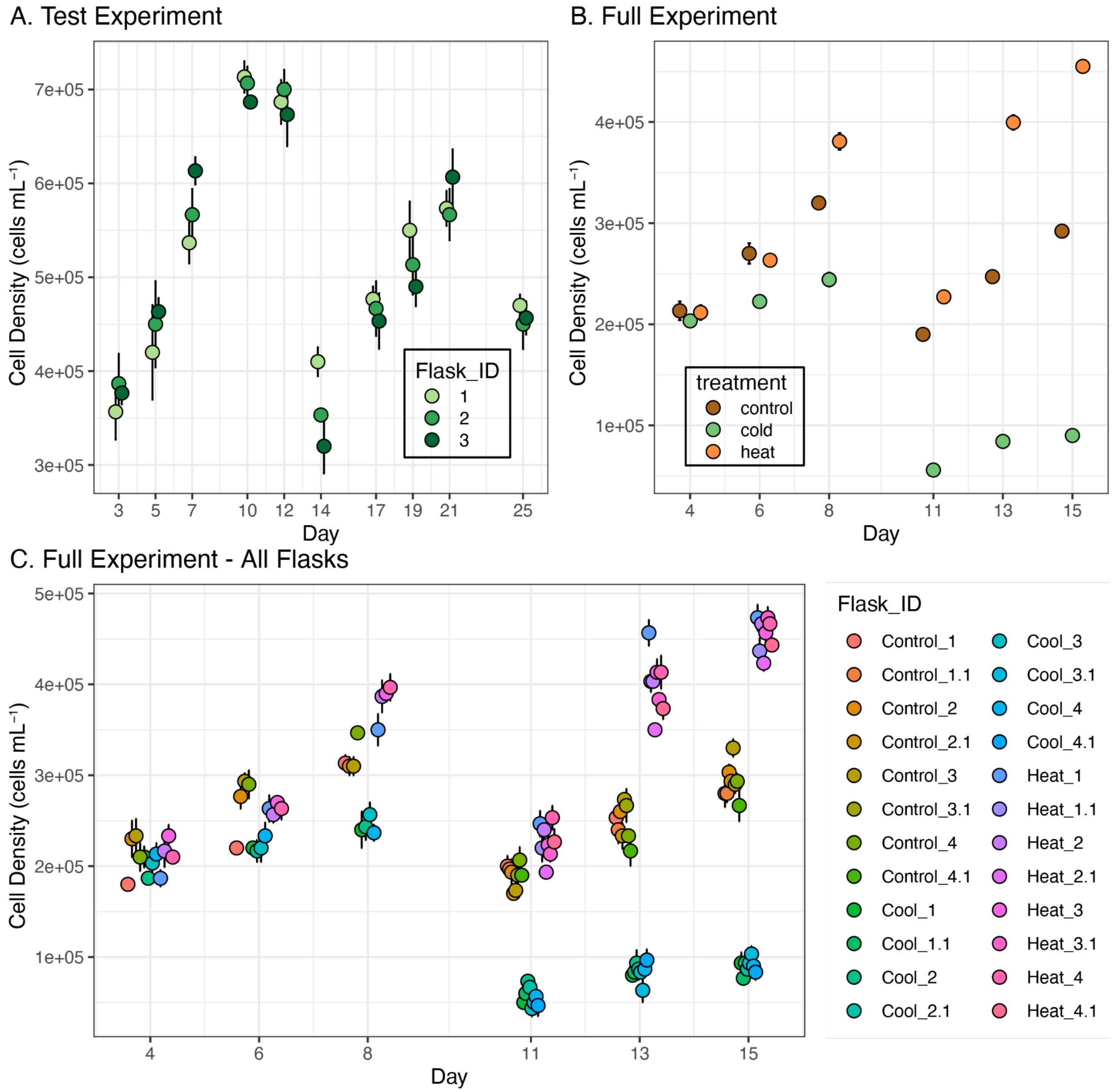


**Figure S3. *Ex hospite Breviolum psygmophilum* cell growth to establish semi-continuous growth methodology. A.** Cell density through time during a test experiment to determine timing of exponential and stationary growth phases of *B. psygmophilum* cultures under control conditions (18°C). The three replicate flasks reached stationary phase on approximately day 12, which informed later transfers to fresh media (**B,C**) to maintain cultures in the exponential growth phase following semi-continuous culturing methodology. **B,C.** Cell density through time during the *ex hospite* symbiont experiment, where transfers to fresh media occurred on day 8. The same data are presented in both **B** and **C**, but data are aggregated across replicate flasks in **B** while data from all individual flasks are shown in **C**.

### Supplemental Tables

**Table S1.** ***Oculina arbuscula*** **and** ***Breviolum psygmophilum*** **holobiont sequencing information.** Sample names (Oculina_ID) indicate genotype, fragment number, temperature treatment (C = control, F = cold challenge, H = heat challenge), and symbiotic state (W = aposymbiotic (white), B = symbiotic (brown)). RawReads indicate the number of reads from the unfiltered fastq file. TrimmedReads indicate the number of reads remaining after filtering. MappedReads are the number of reads aligning one time to the concatenated *O. arbuscula* and *B. psygmophilum* reference transcriptome. HostCounts and SymCounts indicate the *O. arbuscula* and *B. psygmophilum* counts used in DESeq, respectively. Asterisks next to sample ID's indicate samples that were removed, either because they were clones (genotypes O, L, K, A) or because of apparent sequencing failure (genotype A).

| Oculina_ID | RawReads | TrimmedReads | MappedReads | HostCounts | SymCounts |
|---|---|---|---|---|---|
| OA4_C_W* | 713004 | 81024 | 12147 | 18323 | 1089 |
| OA5_F_W* | 3804138 | 957178 | 163218 | 221233 | 40560 |
| OA6_H_W* | 6953917 | 1294706 | 236223 | 344287 | 26010 |
| OC4_F_B | 3129577 | 934048 | 186743 | 288914 | 17844 |
| OC5_H_B | 2795155 | 724676 | 130277 | 180792 | 26135 |
| OC9_C_B | 5259035 | 1208042 | 205301 | 318337 | 22893 |
| OD4_C_B | 3136756 | 951626 | 164800 | 255933 | 21264 |
| OD5_F_B | 6030497 | 1673236 | 334379 | 526111 | 37542 |
| OD6_H_B | 3724669 | 1008182 | 179415 | 271569 | 18113 |
| OE10_F_W | 4196932 | 1245305 | 226276 | 380053 | 16239 |
| OE11_C_W | 3308942 | 909105 | 164122 | 244119 | 26366 |
| OE3_H_W | 4162845 | 1317354 | 232765 | 372206 | 20520 |

| | | | | | |
|---|---|---|---|---|---|
| OF7_C_B | 2097443 | 605880 | 115258 | 156872 | 20246 |
| OF8_F_B | 2937370 | 969666 | 179015 | 297083 | 12948 |
| OF9_H_B | 12787828 | 1776900 | 281666 | 438839 | 25337 |
| OH11_F_W | 2126834 | 693351 | 196295 | 209505 | 9177 |
| OH15_H_W | 8292392 | 2165705 | 126032 | 534770 | 43978 |
| OH1_C_W | 3782080 | 1130597 | 354041 | 303477 | 28026 |
| OI1_C_B | 8138535 | 2203496 | 419720 | 657766 | 47404 |
| OI2_F_B | 4702871 | 1418771 | 271742 | 417661 | 39203 |
| OI3_H_B | 3848414 | 897870 | 148130 | 232085 | 13119 |
| OJ13_C_B | 6221360 | 1833913 | 341585 | 525116 | 38777 |
| OJ14_F_B | 8008008 | 2142125 | 406134 | 640399 | 40860 |
| OJ15_H_B | 2255126 | 724997 | 137730 | 189255 | 32169 |
| OK1_C_W* | 833657 | 321054 | 50955 | 84859 | 4020 |
| OK2_F_W* | 6949550 | 1199628 | 204041 | 354591 | 6864 |
| OK3_H_W* | 1802139 | 660720 | 114659 | 194220 | 6332 |
| OL6_C_B* | 3668299 | 1077721 | 205720 | 304863 | 31519 |
| OL7_F_B* | 4727630 | 1335005 | 256529 | 398154 | 33031 |
| OL8_H_B* | 2466305 | 790985 | 146068 | 216377 | 15962 |
| OM1_C_B | 7295614 | 1938251 | 350216 | 528331 | 37088 |

| | | | | | |
|---|---|---|---|---|---|
| OM2_F_B | 1853944 | 619682 | 121936 | 180069 | 15985 |
| OM3_H_B | 2884074 | 871352 | 178897 | 224827 | 49798 |
| ON4_C_W | 4824787 | 1355099 | 244121 | 404358 | 10913 |
| ON5_F_W | 16246514 | 4022360 | 741401 | 1245818 | 16411 |
| ON6_H_W | 4920306 | 790076 | 115833 | 197072 | 4483 |
| OO1_C_W* | 5739490 | 1659319 | 288043 | 492086 | 7458 |
| OO2_F_W* | 3620699 | 1177521 | 216080 | 364733 | 5697 |
| OP1_C_W* | 5471120 | 1520681 | 260562 | 434079 | 13265 |
| OP2_F_W* | 4596831 | 1275987 | 238375 | 409622 | 8280 |
| OP3_H_W* | 6076638 | 1231158 | 209973 | 320730 | 25153 |
| OQ11_H_W | 8004810 | 1825271 | 156895 | 378358 | 38204 |
| OQ1_C_W | 3568498 | 969113 | 253911 | 269162 | 5799 |
| OQ4_F_W | 9668834 | 2864331 | 556904 | 897646 | 49375 |
| OR7_C_B | 3930279 | 1070544 | 180214 | 284083 | 23315 |
| OR8_F_B | 4983180 | 1287597 | 258211 | 392987 | 27428 |
| OR9_H_B | 4230653 | 676034 | 95841 | 149213 | 12362 |

**Table S2. *Breviolum psygmophilum* in culture (*ex hospite*) sequencing information.** Culture names (Culture_ID) indicate the treatment and replicate number (1-4), and dashes after the replicate number (-1) represent cultures that were split on day 10 to maintain exponential growth using the semi-continuous method. Culture flasks in the cold challenge treatment have four instead of eight replicates because the semi-continuous culture replicates were pooled to obtain sufficient RNA for sequencing. RawReads indicate the number of reads from the unfiltered fastq file. TrimmedReads indicate the number of reads remaining after filtering. Mapped reads are the number of reads aligning one time to the reference transcriptome. Counts indicate the *B. psygmophilum* counts used in DESeq.

| Culture_ID | RawReads | TrimmedReads | MappedReads | Counts |
|---|---|---|---|---|
| Control-1-1 | 8576577 | 3445558 | 1077887 | 1285858 |
| Control-1 | 9322624 | 3355528 | 1056244 | 1259276 |
| Control-2-1 | 8949773 | 3374355 | 1032055 | 1229172 |
| Control-2 | 8092097 | 2439177 | 668422 | 806901 |
| Control-3-1 | 8013989 | 2765579 | 777245 | 931442 |
| Control-3 | 5053389 | 1720274 | 474485 | 571218 |
| Control-4-1 | 6743878 | 2309182 | 610653 | 728048 |
| Control-4 | 6924866 | 2023393 | 505631 | 603137 |
| Cold-1 | 4595701 | 1541637 | 422413 | 484972 |
| Cold-2 | 5492316 | 2006782 | 611776 | 713239 |
| Cold-3 | 6490381 | 2798699 | 968135 | 1113555 |
| Cold-4 | 6515185 | 2871085 | 1015311 | 1184420 |
| Heat-1-1 | 5405332 | 2627373 | 887841 | 1060861 |
| Heat-1 | 7322437 | 2939714 | 975190 | 1144023 |
| Heat-2-1 | 7897004 | 2661411 | 830809 | 983978 |

| | | | | |
|---|---|---|---|---|
| Heat-2 | 6252561 | 2847025 | 983662 | 1154335 |
| Heat-3-1 | 6653011 | 2909299 | 980661 | 1154315 |
| Heat-3 | 6055500 | 1875336 | 512730 | 606975 |
| Heat-4-1 | 7074897 | 2946953 | 1006165 | 1186170 |
| Heat-4 | 6879771 | 2115655 | 580844 | 684762 |

**Table S3. Ortholog counts.** Counts indicate the ortholog counts for *Oculina arbuscula* (.host Sample Name) and *Breviolum psygmophilum* (.sym Sample Name) counts used in DESeq (clones removed).

| Sample Name | Counts |
|---|---|
| OC4_F_B.host | 30064 |
| OC5_H_B.host | 15342 |
| OC9_C_B.host | 28894 |
| OD4_C_B.host | 22375 |
| OD5_F_B.host | 54748 |
| OD6_H_B.host | 22043 |
| OE10_F_W.host | 40217 |
| OE11_C_W.host | 22239 |
| OE3_H_W.host | 31446 |
| OF7_C_B.host | 14277 |
| OF8_F_B.host | 32754 |
| OF9_H_B.host | 38000 |
| OH11_F_W.host | 22030 |
| OH15_H_W.host | 45246 |
| OH1_C_W.host | 26822 |
| OI1_C_B.host | 56139 |
| OI2_F_B.host | 44721 |
| OI3_H_B.host | 19673 |
| OJ13_C_B.host | 44670 |

| | |
|---|---|
| OJ14_F_B.host | 66362 |
| OJ15_H_B.host | 15506 |
| OM1_C_B.host | 42803 |
| OM2_F_B.host | 18597 |
| OM3_H_B.host | 18152 |
| ON4_C_W.host | 36191 |
| ON5_F_W.host | 124340 |
| ON6_H_W.host | 17108 |
| OQ11_H_W.host | 32114 |
| OQ1_C_W.host | 23850 |
| OQ4_F_W.host | 93015 |
| OR7_C_B.host | 25069 |
| OR8_F_B.host | 41554 |
| OR9_H_B.host | 12651 |
| OC4_F_B.sym | 1970 |
| OC5_H_B.sym | 2060 |
| OC9_C_B.sym | 2313 |
| OD4_C_B.sym | 2018 |
| OD5_F_B.sym | 3693 |
| OD6_H_B.sym | 1630 |
| OF7_C_B.sym | 1757 |

| | |
|---|---|
| OF8_F_B.sym | 1773 |
| OF9_H_B.sym | 2349 |
| OI1_C_B.sym | 3632 |
| OI2_F_B.sym | 3811 |
| OI3_H_B.sym | 1261 |
| OJ13_C_B.sym | 3117 |
| OJ14_F_B.sym | 4169 |
| OJ15_H_B.sym | 2180 |
| OM1_C_B.sym | 3073 |
| OM2_F_B.sym | 1621 |
| OM3_H_B.sym | 3627 |
| OR7_C_B.sym | 1815 |
| OR8_F_B.sym | 2733 |
| OR9_H_B.sym | 1059 |
